# Survey of commercial antibodies targeting Y chromosome-encoded genes

**DOI:** 10.1101/2023.07.26.550552

**Authors:** Bradley D. Gelfand, Dionne A. Argyle, Joseph J. Olivieri, Jayakrishna Ambati

**Affiliations:** Center for Advanced Vision Science, University of Virginia School of Medicine, Charlottesville, VA, USA; Department of Ophthalmology, University of Virginia School of Medicine, Charlottesville, VA, USA; Department of Biomedical Engineering, University of Virginia School of Medicine, Charlottesville, VA, USA; Department of Pathology, University of Virginia School of Medicine, Charlottesville, VA; Department of Microbiology, Immunology, and Cancer Biology, University of Virginia School of Medicine, Charlottesville, VA, USA

## Abstract

Commercial antibodies are both a mainstay of biomedical research and subject to increasingly widespread scrutiny for their specificity. We evaluated the specificity of commercially available research grade antibodies by investigating those whose antigen targets are encoded by the Y chromosome. We assessed vendor-provided validation data such as immunoblotting and identified 65 commercial research grade antibodies that exhibited positive reactivity in female-derived tissues and/or cell lines. A thorough analysis of 30 commercial antibodies marketed as targeting the Y chromosome-encoded protein DDX3Y found that 16 (53%) provided no marketing data on specificity, 9 (30%) provided positive data in female or likely female materials, 4 (13%) provided only positive data in male materials, and 1 (3%) provided positive data in male materials and negative data in female materials. Together, these findings suggest that many commercial antibodies may be unsuitable to distinguish between related X and Y chromosome-encoded gene pairs and should instill caution in researchers using these tools to investigate sex chromosome-encoded proteins.

## Introduction

Although immunoassays are an indispensable tool for scientific research, the antibody specificity has been recognized as a major challenge to the rigor and reproducibility of research findings. A 2016 proposal published by the International Working Group for Antibody Validation identified five pillars of antibody validation.^1^ Among these is genetic validation, in which “The expression of the target protein is eliminated or significantly reduced by genome editing or RNA interference”.

Y chromosome-encoded genes present unique opportunities and challenges to validate antibodies on this genetic principle. Fortunately, readily available female-derived cells and tissues can serve as a target-negative source material, which is far more convenient than typical sources of genetic validation which require knockout or knockdown approaches to a target gene. However, an additional challenge for specificity of these antibodies is that many Y chromosome proteins have “gametologs”, or highly homologous genes encoded on the X chromosome. As gametologs can share over 90% amino acid identity, these protein targets present unique specificity challenges. However, this obstacle has not impeded commercial antibody suppliers who market hundreds of antibodies with purported specificity for Y chromosome-encoded genes. Here we present findings from an analysis of the extent to which Y chromosome gene-targeted commercial antibodies recognize female derived materials using data provided in their marketing materials. We identified 65 antibodies purporting to recognize Y chromosome-encoded genes with supplier provided data showing reactivity in female tissues.

## Methods

Antibodies purporting to recognize Y chromosome targeted genes were identified by using the Antibody Search tool from Biocompare,^2^ accessed between September 2022 and April 2023. Searches were conducted for the Y chromosome gene name and occasionally for commonly used aliases, which were identified via National Library of Medicine Gene resource. The searches were limited to those with human reactivity based on the Biocompare search tool settings.

We reviewed the company-supplied validation data and attempted to identify the sex of the source material. The chromosomal sex of cell lines was identified using Expasy’s Cellosaurus database.^3^ The sex was unambiguously identified in all but one cell line. One antibody purporting to target HSFY1 listed COLO205 cells as the source. As both male (CVCL_0218) and female (CVCL_F402) listings for COLO205 cells were found in Cellosaurus, this antibody was excluded from our analysis. Best attempts were made to ensure the same antibodies supplied by different companies were not double counted by comparing available validation data and other details (clone, antigen, source, application, etc.).

## Results

**Table 1** lists 65 antibodies purporting to target a Y chromosome-encoded gene with company-supplied marketing demonstrating immunoreactivity in female-derived tissues. For one example, an antibody targeting Sex determining region chromosome Y (SRY) marketed by MyBioSource (catalog # MBS8513980) presents validation data in HeLa cells, which is a cervical cancer cell line with no Y chromosomes.^4^

**Table 1.**
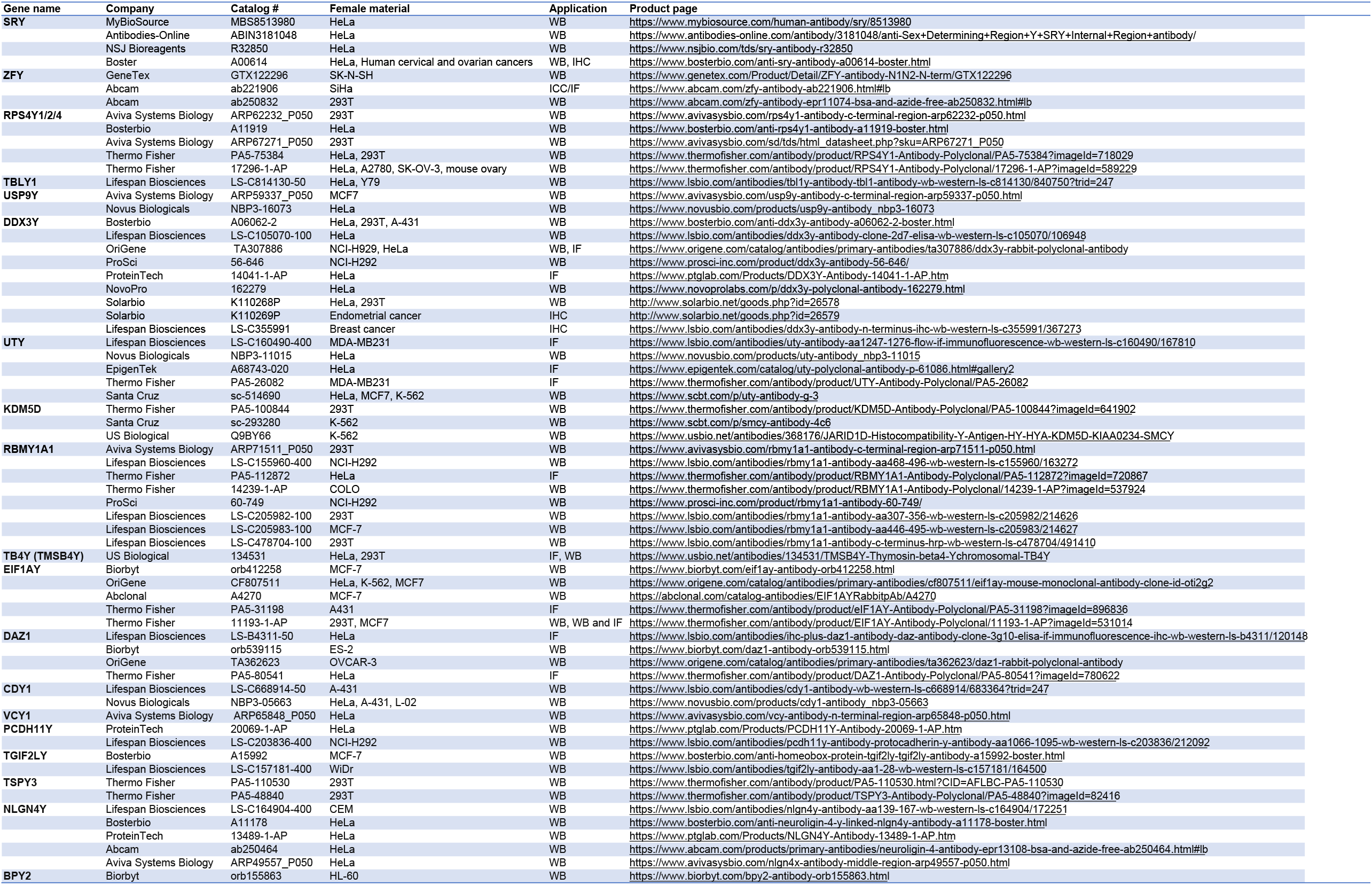
Commercial antibodies against Y chromosome encoded proteins with positive immunoreactivity in female materials organized by gene name. Abbreviations: WB = western blotting, IHC = immunohistochemistry, ICC = immunocytochemistry, IF = immunofluorescence.

Among these antibodies, frequently used female-derived cell lines were HeLa, 30/65 (46%), HEK293T, female human embryonic kidney cells used in 14 (22%), and MCF-7 breast cancer cells used in 7 (11%). One antibody, a rabbit polyclonal raised against the “N terminus” of DDX3Y (LS Biosciences, cat # LS-C355991) presented positive immunohistochemistry in human breast cancer tissue. While not definitively a Y-chromosome absent tissue, we included this as a likely female positive tissue, based on the prevalence of breast cancer in females compared to males being roughly 99-to-1 in the United States.^5^

Among 65 antibodies, we noted just two that had disclaimers warning that the antibody may cross-react with homologous X chromosome-encoded proteins.

Results of a detailed analysis of all commercially available antibodies targeting DEAD-box helicase 3 Y-linked (DDX3Y) is presented in **Table 2**. DDX3Y is a gametolog of the X chromosome-encoded gene DDX3X, with ∼92% homology between the proteins. We identified 30 antibodies purporting to target DDX3Y and assigned them into four categories based on marketing validation data criteria as follows:

**Table 2.**
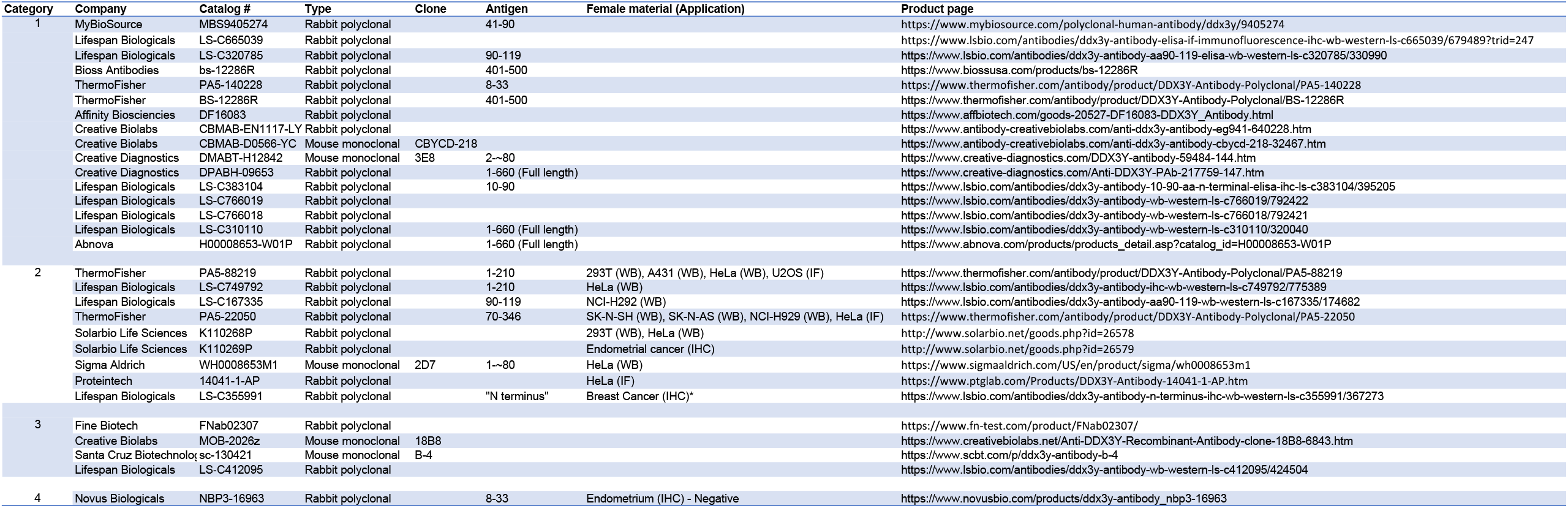
Commercial antibodies targeting Y chromosome encoded DDX3Y. Clone, antigen, and female positive results as provided by company marketing materials. *Positive reactivity in human breast cancer tissue is presumed to be Y chromosome absent, and therefore indicative of non-specific staining.

1. No validation data provided (16/30, 56%)
2. Validation data provided indicating positive signal in female or likely female tissue (see discussion below) with or without positive data in male tissue (9/30, 30%)
3. Validation data provided indicating positive signal in male or likely male tissue, but no data on female tissue (4/30, 13%)
4. Validation data provided indicating positive signal in male or likely male tissue, and affirmatively negative data in female tissue (1/30, 3%)

## Discussion

We provide evidence of widespread off-target antigen recognition in commercial antibodies purporting to recognize Y chromosome-encoded proteins. Some important caveats should be noted.

First, many antibodies provided no primary data on female tissues. For example, 20/30 (67%) of DDX3Y antibodies provided no data in female tissues. Therefore, it seems likely that the 65 antibodies listed in **Table 1** are a significant underrepresentation of Y chromosome-targeted antibodies lacking specificity.

Second, this analysis assumes that the identities of the listed cell types provided in marketing materials are accurate and not subject to cell line contamination, conceivably with Y chromosome-containing cells. Cell line purity and identity is itself a major challenge in biomedical research.

Third, in the case of human tissues such as endometrium and breast cancer, it is conceivable that positive immunoreactivity from Y chromosome-encoded proteins could represent true staining of microchimerism, in which an allogeneic cell population resides within a host. We think this is unlikely because even in the extreme case where every Y chromosome-containing cell expresses the antigen, one would expect a true positive staining pattern to be restricted the small number of allogenic cells as in other in situ hybridization staining analyses of microchimeric tissues,^6^ rather than widespread staining as is reported in antibody marketing materials.

In summary, many commercial antibodies targeting Y chromosome-encoded proteins are not validated for use in sex-specific applications. Researchers using these tools are encouraged to validate their reagents in tissues lacking a Y chromosome and should be cautious when interpreting findings regarding antigens encoded by the Y chromosome. We also urge commercial antibody suppliers to provide better warning to consumers about the lack of validated specificity among Y chromosome-targeted antibodies.

